# DamageProfiler: Fast damage pattern calculation for ancient DNA

**DOI:** 10.1101/2020.10.01.322206

**Authors:** Judith Neukamm, Alexander Peltzer, Kay Nieselt

## Abstract

In ancient DNA research, the authentication of ancient samples based on specific features remains a crucial step in data analysis. Because of this central importance, researchers lacking deeper programming knowledge should be able to run a basic damage authentication analysis. Such software should be user-friendly and easy to integrate into an analysis pipeline. Here, we present DamageProfiler, a Java based, stand-alone software to determine damage patterns in ancient DNA. The results are provided in various file formats and plots for further processing. DamageProfiler has an intuitive graphical as well as command line interface that allows the tool to be easily embedded into an analysis pipeline.

## 1 Introduction

The field of ancient DNA (aDNA) further develops with every passing year and current research spans diverse topics including the analysis of various taxa, from ancient plant and animal species (Jaenicke-Després *et al*., 2003; Debruyne ; *et al.*, 2008), to human microbiological communities from oral or gut microbiomes (Warinner, 2016; Tito *et al.*, 2012). One central analysis step is the authentication of the ancient origin of the DNA. Several characteristics have been identified that describe postmortem DNA modifications. These can be used to differentiate between authentic ancient DNA and modern contamination and are typically summarized as (a) short fragment length (Meyer *et al.*, 2016); (b) preferential fragmentation of ancient DNA at purine bases (Briggs *et al.*, 2007); and (c) increased Cytosine (C) to Thymine (T) base misincorporations towards the five prime (5’) end and (d) complementary Guanine (G) to Adenine (A) base misincorporations towards the three prime (3’) end of the fragment due to enhanced cytosine deamination in single-stranded 5’-overhanging ends (Briggs *et al.*, 2007). Existing software such as mapDamage2 (Jónsson *et al.*, 2013) calculates the nucleotide misincorporation and fragmentation patterns from next-generation sequencing (NGS) reads mapped against a reference genome and provides graphical and text-based representations, as well as further statistical estimates. However, its usability is hindered by the exclusive use as a command line tool and long runtimes for larger samples sets. For ancient metagenomic samples, HOPS (Hübler *et al.*, 2019) has recently been published. This tool offers a fully automatized bacterial screening pipeline for ancient DNA sequences that provides detailed information on species identification and authenticity. However, HOPS includes the metagenomic mapping software MALT (Vågene *et al.*, 2018) and is limited to its output format.

Here, we present DamageProfiler, a fast and user-friendly software to investigate damage patterns in high-throughput sequencing data. DamageProfiler aims for high usability and fast runtime to be applicable to large data sets. The results are given as intuitive, well-described figures and text-based representation in various file formats, including json format. Moreover, the visualizations can be explored interactively in the graphical user interface. The various output formats and the improvement of runtimes makes it suitable for integrated use in analysis pipelines.

## 2 Results

DamageProfiler is a fast, interactive, and easy-to-use program to investigate the damage patterns of high-throughput sequencing data. DamageProfiler is compatible with newer file formats such as CRAM, and also legacy file formats such as SAM or BAM. In addition to several statistics, DamageProfiler generates a damage plot, which is calculated by determining the frequency of C to T base misincorporations per position of all fragments that map to the reference genome, the length distribution of all mapped reads, and the edit distance between the read and the reference both in text and image formats (figure 1).

**Figure 1:**
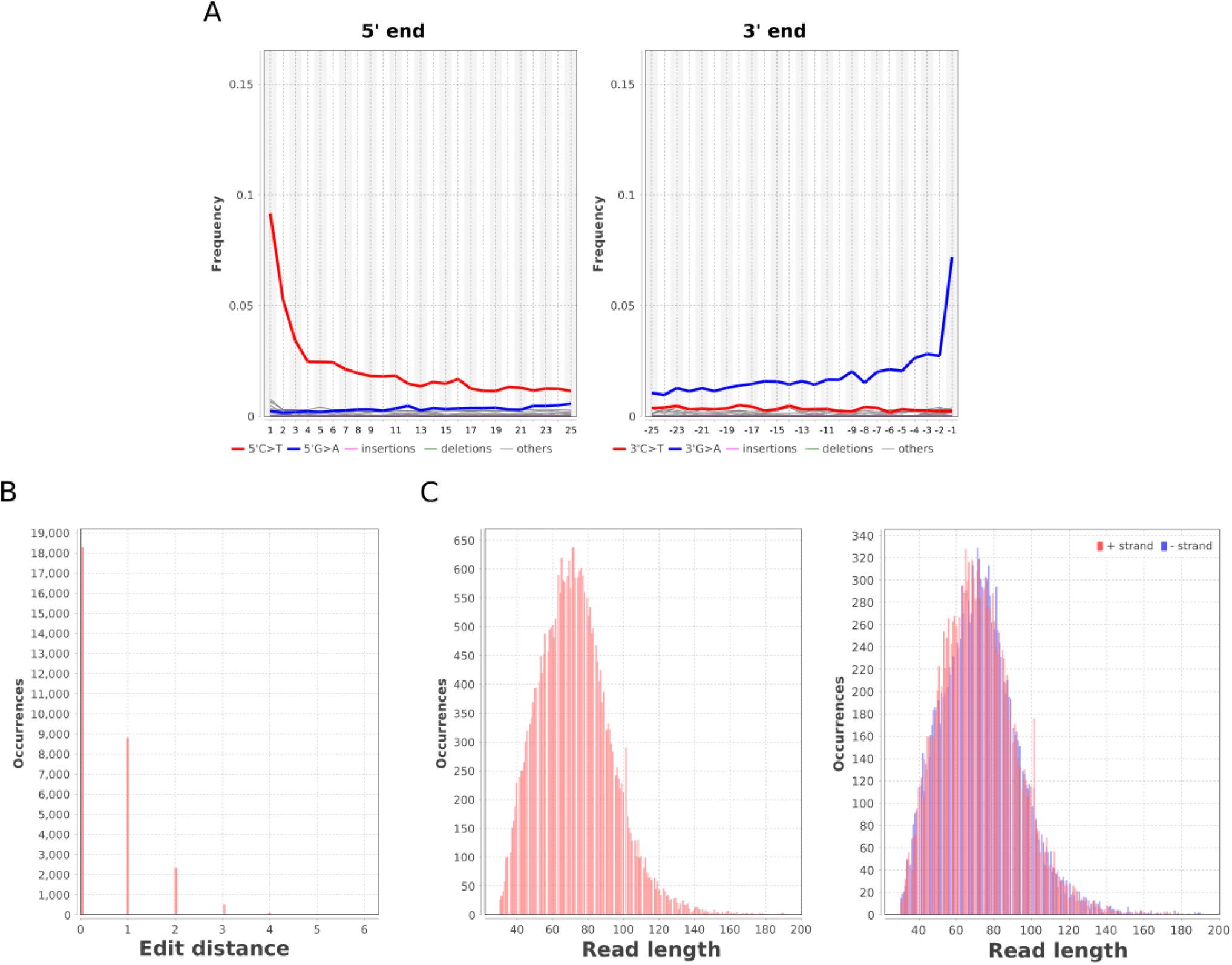
DamageProfiler on mapping with BWA-aln. Results of DamageProfiler based on the mapping file of sample JK2134 created using BWA-aln. As reference, the human mitochondrial genome (NC_012920.1) was used. In total, 30,064 reads could be mapped. The (A) frequency of base misincorporation at the ends of the fragment, the (B) edit distance between the reads and the reference, and the (C) read length distribution for each strand individually and the combination are shown.

Contrasting to the double-stranded aDNA library preparation protocol (Briggs and Heyn, 2012), the single-stranded library protocol (Gansauge *et al.*, 2017) results in reads having only C to T base misincorporations at both the 5’ and 3’ ends and thus renders the visualization of the G to A base misincorporations superfluous. Subsequently, for single-stranded protocols, DamageProfiler provides a specific option to compute and visualize only C to T base misincorporations.

DamageProfiler is also applicable to metagenomic mapping files, i.e. mapping files that contain mapping information for multiple reference files. Based on a user-specified list of reference IDs, a detailed output for each species and a general summary file are generated.

DamageProfiler runs platform-independent using the Java virtual machine (JVM) version 11 and higher and provides both a graphical and command-line interface. The graphical user interface (supplementary figure S1) enables easy configuration of all parameters and offers an interactive, visual exploration of the results. All utilized parameters and steps performed are listed in a log file, which helps to track each step for reproducible results. In addition to comprehensive documentation (https://damageprofiler.readthedocs.io), a help page is available directly within the interface to provide more details about all parameters.

DamageProfiler is developed as stand-alone software, but can also be integrated into analysis pipelines. Currently, DamageProfiler is already available within the EAGER pipeline (Peltzer *et* al., 2016), and is the default damage computation tool in nf-core/eager (Fellows Yates *et al.*, 2020; Ewels *et al.*, 2020).

DamageProfiler accepts BAM, SAM, and CRAM formatted files and provides various output formats, such as SVG images, text, and JSON files, for further processing of the results. The JSON output can, for example, be used to summarize results with direct integration in MultiQC (Ewels *et al.*, 2016).

To evaluate the accuracy of the results, DamageProfiler was tested on a simulated dataset (supplementary note 1, figure S2). We could show that the results are identical to the expected damage patterns, which proves the reliability of DamageProfiler.

While various mapping software are used in the ancient DNA field, DamageProfiler was successfully tested on the most common ones, such as Bowtie2 (Langmead and Salzberg, 2012), bwa-aln (Li and Durbin, 2009), bwa mem (Li, 2013), and MiniMap2 (Li, 2018) (supplementary note 2, figure S3).

Moreover, we demonstrated the application of DamageProfiler on a metagenomic mapping file (supplementary note 3, figure S4). Based on a given set of species of interest, DamageProfiler calculates the damage patterns for all species individually and also generates a summary file containing all species.

DamageProfiler also improves the runtime of damage pattern calculations compared to mapDamage2 (Jónsson *et al.*, 2013). As mapDamage and DamageProfiler provide different additional features, we only compared the runtime of accessing the differences between the actual mapping process and the output generation. The runtime was tested for four files of different sizes and reference genomes and it could be shown that DamageProfiler runs 4.4 times faster on average than mapDamage2 (supplementary note 4, table S1). Moreover, we observed that the runtime improves with increasing size of the mapping file (supplementary note 4, figure S5).

## 3 Discussion

We demonstrated the reliability of the results of DamageProfiler and showed that our tool is on average 4.4 times faster than the existing software mapDamage2. An increase in speed can be specifically observed for larger mapping files, and could even be improved by a multithreaded implementation.

In summary, DamageProfiler is a fast and user-friendly tool for coherent calculation of damage patterns for mapped NGS reads from a variety of mapping tools. Furthermore, DamageProfiler can be applied to single- or multi-reference mapping files.

## Acknowledgments

We would like to thank Abagail M. Breidenstein for proofreading the manuscript. We also thank Maxime Borry, Alexander Hübner, James A. Fellows Yates, and Simon Heumos for bug reports, feature suggestions, and contributions.

## Data Availability

All of the source code is freely available on GitHub (https://github.com/Integrative-Transcriptomics/DamageProfiler). Detailed documentation can be found online: https://damageprofiler.readthedocs.io.

## References

Briggs,A.W. et al. (2007) Patterns of damage in genomic DNA sequences from a Neandertal. Proc. Natl. Acad. Sci. U. S. A., 104, 14616–14621.

Briggs,A.W. and Heyn,P. (2012) Preparation of Next-Generation Sequencing Libraries from Damaged DNA. In, Shapiro,B. and Hofreiter,M. (eds), Ancient DNA: Methods and Protocols. Humana Press, Totowa, NJ, pp. 143–154.

Debruyne,R. et al. (2008) Out of America: ancient DNA evidence for a new world origin of late quaternary woolly mammoths. Curr. Biol., 18, 1320–1326.

Ewels,P. et al. (2016) MultiQC: summarize analysis results for multiple tools and samples in a single report. Bioinformatics, 32, 3047–3048.

Ewels,P.A. et al. (2020) The nf-core framework for community-curated bioinformatics pipelines. Nat. Biotechnol., 38, 276–278.

Fellows Yates, J.A. et al. (2020) Reproducible, portable, and efficient ancient genome reconstruction with nf-core/eager. bioRxiv, 2020.06.11.145615.

Gansauge,M.-T. et al. (2017) Single-stranded DNA library preparation from highly degraded DNA using T4 DNA ligase. Nucleic Acids Res., 45, e79.

Hübler,R. et al. (2019) HOPS: automated detection and authentication of pathogen DNA in archaeological remains. Genome Biol., 20, 280.

Jaenicke-Després,V. et al. (2003) Early allelic selection in maize as revealed by ancient DNA. Science, 302, 1206–1208.

Jónsson,H. et al. (2013) mapDamage2.0: fast approximate Bayesian estimates of ancient DNA damage parameters. Bioinformatics, 29, 1682–1684.

Langmead,B. and Salzberg,S.L. (2012) Fast gapped-read alignment with Bowtie 2. Nat. Methods, 9, 357–359.

Li,H. (2013) Aligning sequence reads, clone sequences and assembly contigs with BWA-MEM. arXiv [q-bio.GN].

Li, H. (2018) Minimap2: pairwise alignment for nucleotide sequences. Bioinformatics, 34, 3094–3100.

Li, H. and Durbin, R. (2009) Fast and accurate short read alignment with Burrows-Wheeler transform. Bioinformatics, 25, 1754–1760.

Meyer, M. et al. (2016) Nuclear DNA sequences from the Middle Pleistocene Sima de los Huesos hominins. Nature, 531, 504–507.

Peltzer, A. et al. (2016) EAGER: efficient ancient genome reconstruction. Genome Biol., 17, 60.

Tito, R.Y. et al. (2012) Insights from characterizing extinct human gut microbiomes. PLoS One, 7, e51146.

VågeneÅ J. et al. (2018) Salmonella enterica genomes from victims of a major sixteenth-century epidemic in Mexico. Nat Ecol Evol, 2, 520–528.

Warinner, C. (2016) Dental Calculus and the Evolution of the Human Oral Microbiome. J. Calif. Dent. Assoc., 44, 411–420.

