## Supplementary Information for "DamageProfiler: Fast damage pattern calculation for ancient DNA"

### Supplementary Note 1: Evaluation of correctness

We used a simulated dataset to evaluate the correctness of DamageProfiler. Gargammel (Renaud *et al.*, 2017) was used to create an Illumina HiSeq 2500 dataset with 1 million fragments and a fragment length of 55 bp. A deamination pattern with 18%, 11% and 7% C to T misincorporation frequency at the first, second and third position at the 5' end was simulated. The frequency of G to A misincorporations at the 3' end was set identically. The simulated reads were processed using EAGER (Peltzer *et al.*, 2016). The reads were adapter trimmed and read-pairs merged based on a minimum overlap of ten bases with AdapterRemoval v2.2.1a (Schubert *et al.*, 2016) and subsequently aligned to the reference genome provided by gargammel using BWA aln v0.7.17 (Li and Durbin, 2009) with a seed length of 1024, a minimum quality score of 25 and a maximum edit distance of n=0.01. Duplicates were removed with MarkDuplicates v2.15.0 (http://broadinstitute.github.io/picard/). Next, DamageProfiler was applied on the resulting mapping file, showing the expected damage patterns (Figure S2A). Lastly, the results were compared with those obtained with mapDamage2 (Jónsson *et al.*, 2013) (Figure S2B), yielding identical results.

### Supplementary Note 2: Mapping software comparison and compatibility

To demonstrate the compatibility with different mapping software, DamageProfiler was run on mapping files generated using four different mapping methods: Bowtie2 version 2.3.4.1 (Quadrato *et al.*, 2017), BWA aln version 0.7.17 (Li and Durbin, 2009), BWA mem version 0.7.17 (Li, 2013) and MiniMap2 (Li, 2018). We used previously published sequencing data of an Egyptian mummy dated to 776-569 cal BC (Schuenemann *et al.*, 2017) and the human mitochondrial genome as a reference (NC_012920.1). First, the raw data were adapter-clipped and paired-end reads merged as described above. Second, four different mapping software were used to align the sequencing reads against the reference genome. For BWA aln, we used the parameters which are best for ancient DNA (Schubert *et al.*, 2012; Kircher, 2012). Bowtie2 was run using the parameters for ancient DNA suggested by EAGER (Peltzer *et al.*, 2016). MiniMap2 was run with the option to perform a mapping for short reads ('-sr'). BWA-mem was evaluated with the default parameters. If available, the MD tag was calculated as well. This tag contains information about the mismatching positions of the record and renders the reference file superfluous when calculating the corresponding reference of a record.

**Mapping commands:**

**Bowtie:**

```
bowtie2 -x reference.fasta -U input.fq.gz -S output.sam --end-to-end --very-sensitive -p 8
```

**BWA aln:**

```
bwa aln -t 8 -n 0.01 -l 1024 -o 2 -f output.sai input.fq.gz
bwa samse -r reference.fasta output.sai input.fq.gz -f output.sam
```

**BWA-mem:**

```
bwa mem -t 8 reference.fasta input.fq.gz > output.sam
```

**MiniMap2:**

```
minimap2 --MD --eqx -t 8 -ax sr reference.fasta input.fq.gz > aln.sam
```

Next, the mapping files were converted to BAM files, unmapped records were excluded, only read operations with a mapping quality of more than 25 were kept and duplicates were removed. DamageProfiler was executed on the resulting mapping file:

```
java -jar DamageProfiler-1.0.jar -i input.bam -o output_folder -title ‘JK2134 (mapper)’
-yaxis_dp_max 0.4
```

The similarity of the results is shown in figure S3.

### Supplementary Note 3: Metagenomic mapping

DamageProfiler was applied to shotgun sequencing data of a human calculus sample (SRR957739) (Warinner *et al.*, 2014). The file was produced using MALT (Herbig *et al.*, 2016; Vågene *et al.*, 2018) using all complete bacterial, viral and archaeal genomes in GenBank (Benson *et al.*, 2013) as a reference (version May 2018). MALT was executed with the following mapping parameters: Only reads with a minimum 85% identity (--minPercentIdentity) were considered as a possible match to the reference. The minimum support parameter (--minSupport) was set to 5, i.e. only nodes with minimum support of five reads are kept. BlastN mode and SemiGlobal alignment were applied and a top percent value (--topPercent) of 1 was set. All other parameters were set to default. The damage patterns were calculated for a given list of species, containing the top four species identified: Actinomyces sp. oral taxon 414 (NZ_CP012590.1), *Streptococcus cristatus* (NC_021175.1), *Streptococcus sanguinis* (NC_009009.1), and Anaerolineaceae bacterium oral taxon 439 (NZ_CP017039.1), as well as some typical oral bacteria: *S. mutans* (NC_004350.2), *F. alocis* (NC_016630.1), *O. uli* (NC_014363.1) and *T. forsythia* (NC_016610.1), and other typical oral bacteria. The detailed results for each species are stored in the output folder and visualized in the GUI if it was used. Moreover, a summary PDF is created, providing an overview of the results for all species (Figure S4).

### Supplementary Note 4: Runtime comparison

The runtime of DamageProfiler and mapDamage2 (Jónsson *et al.*, 2013) was compared based on four bam files of different size, hosts species and reference genomes (Table S2). The mapping was performed using EAGER as described above, choosing parameters that are optimal for ancient DNA (Schubert *et al.*, 2012; Kircher, 2012). The runtime was measured on Ubuntu 18.04 LTS with 8 cores and 16GB RAM. We only compared the runtime required for the functionality that is shared by mapDamage2 and DamageProfiler, i.e. reading the bam file, and calculating and plotting the damage patterns.

### Supplementary Figures

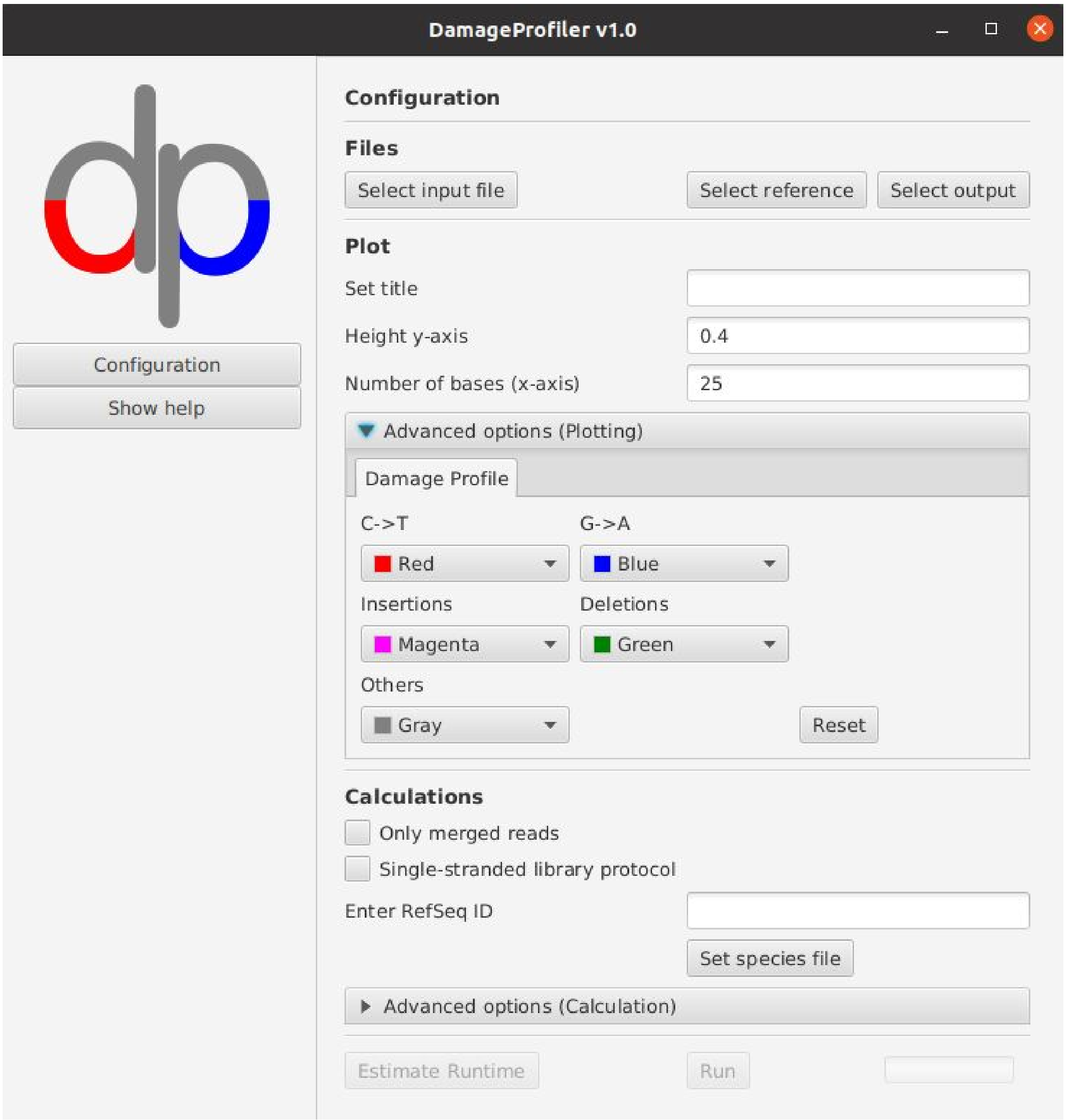


**Figure S1:** Graphical user interface for configuration of a run and visualization of the results.

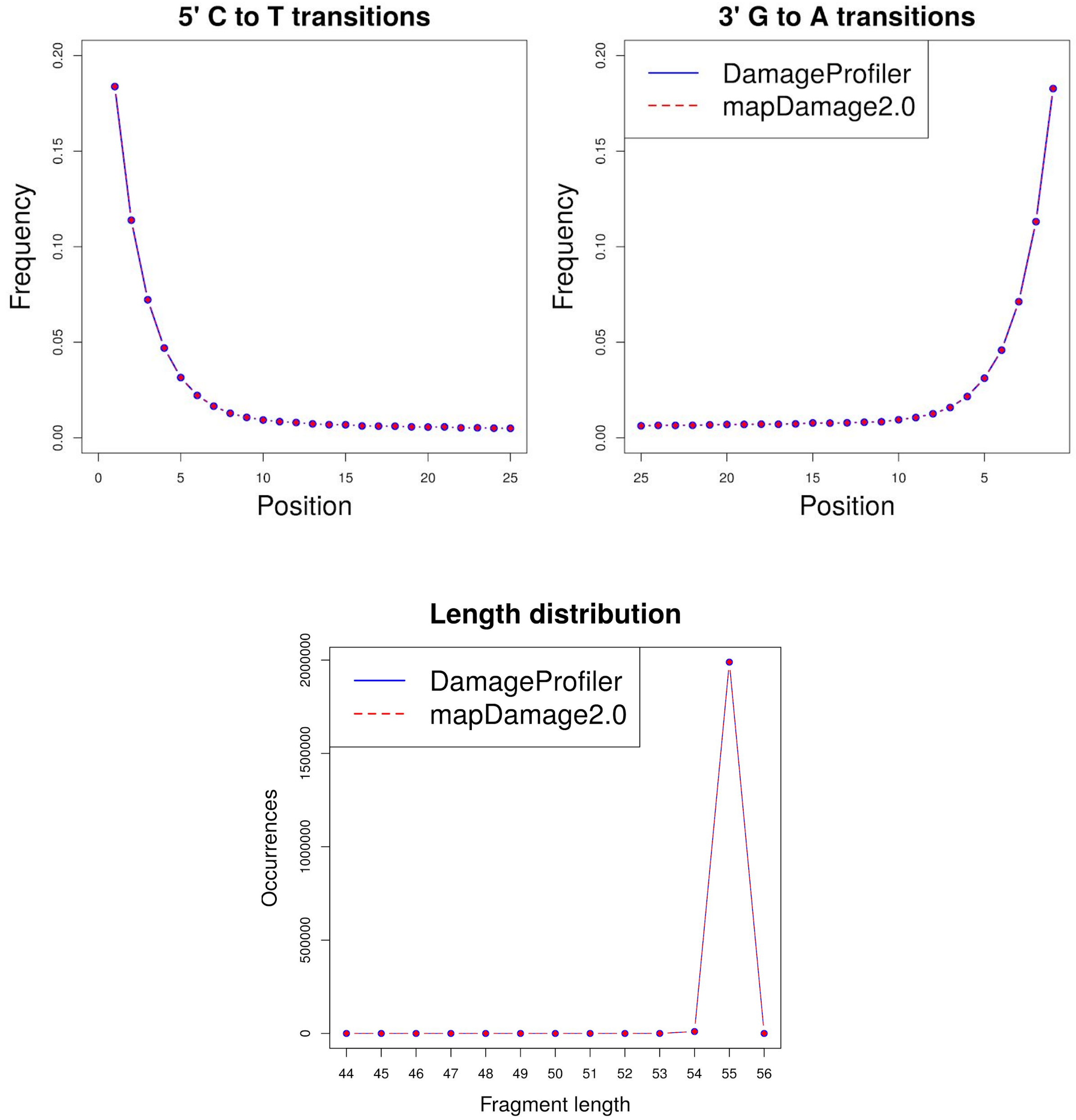


**Figure S2: Damage patterns of simulated data set**. The damage profile and length distribution of the simulated data set calculated with DamageProfiler and mapDamage2 resulting in identical damage patterns.

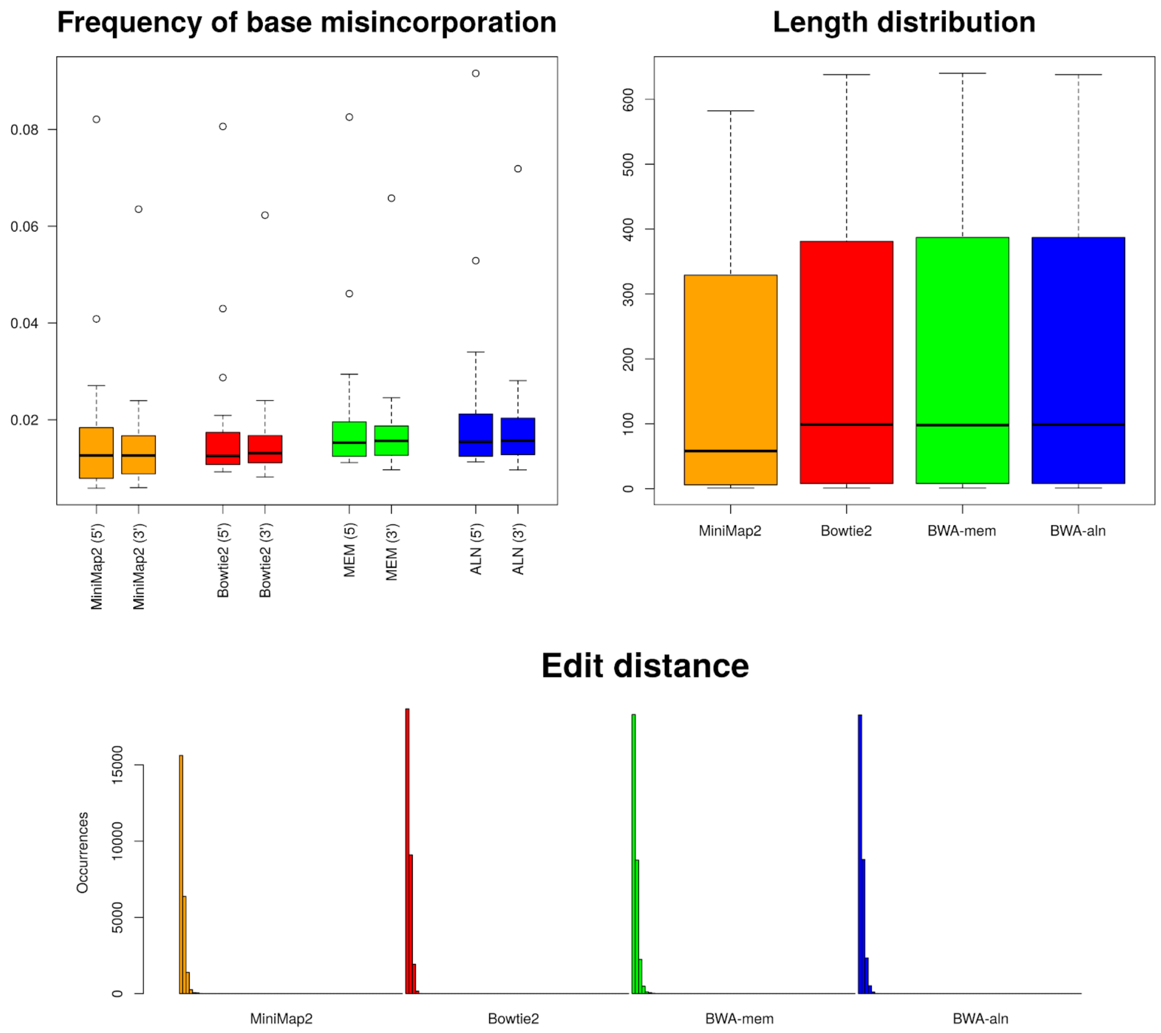


**Figure S3: Comparison of different mappers.** DamageProfiler applied to mapping results of four different mapping software. The base misincorporation, length distribution and edit distance are compared.

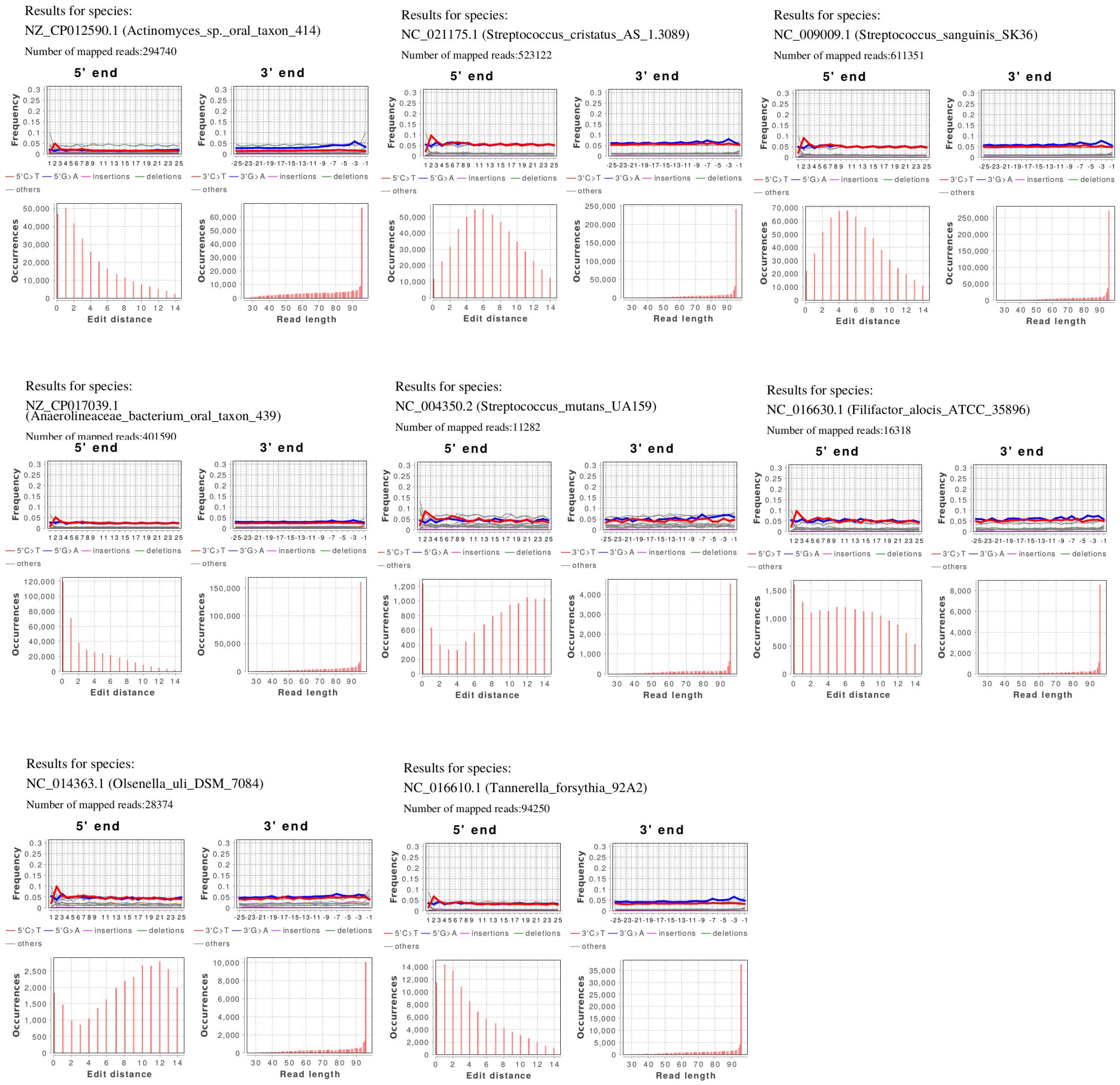


**Figure S4: DamageProfiler on metagenomic mapping file.** Summarized results of DamageProfiler executed on a metagenomic mapping file from a calculus sample and assembled list of oral species (Actinomyces sp. oral taxon 414, *S. cristatus*, *S. sanguinis*, Anaerolineaceae bacterium oral taxon 439, *S. mutans*, *O. uli*, *F. alicos*, *T. forsythia*, *T. denticola* and *P. gingivalis*). A summary of the damage patterns for each species is represented per page. For more detailed results, the complete output of each species is saved in individual folders.

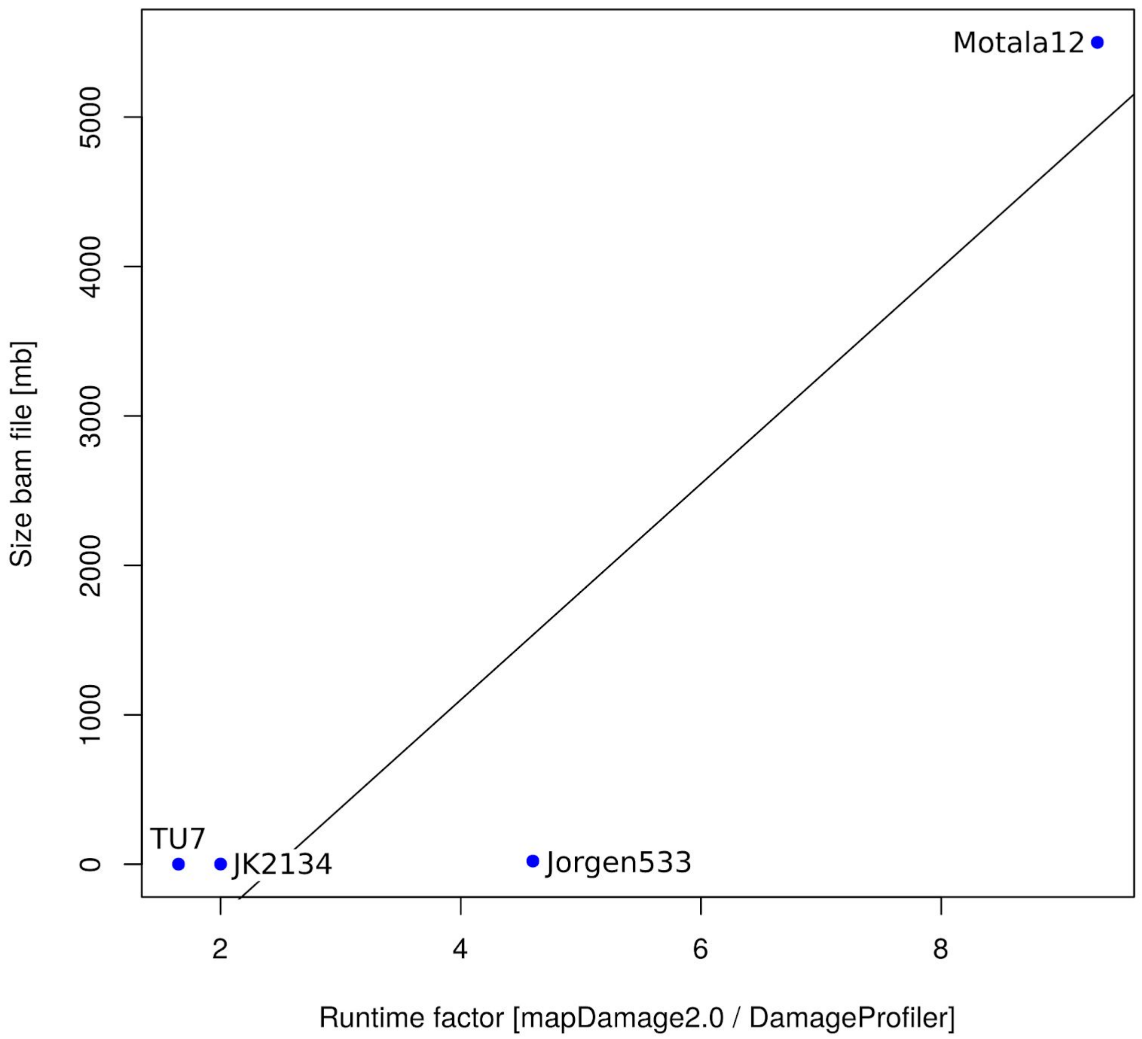


**Figure S5: Runtime factor vs. file size for four different samples.** The runtime factor is shown on the x-axis, which is calculated by dividing the runtime of mapDamage2 by the runtime of DamageProfiler. The runtime represents the average time needed based on ten runs. The y-axis represents the size of the bam file for each sample in megabytes [mb].

### Supplementary Tables

**Table S1: Runtime comparison between mapDamage2 and DamageProfiler.**

| **Sample** | **Publication** | **Ref. genome** | **Size bam file [mb]** | **mapDamage2 Runtime [s]** | **DamageProfiler Runtime [s]** | **Factor** |
|---|---|---|---|---|---|---|
| TU7 | (Gretzinger *et al.*, 2019) | NC_011112.1 | 0.7 | 3.3 | 2.0 | 1.7 |
| JK2134 | (Schuenemann *et al.*, 2017) | NC_012920.1 | 1.5 | 4.2 | 2.1 | 2.0 |
| Jorgen533 | (Schuenemann *et al.*, 2018) | NC_002677.1 | 21.6 | 28.3 | 6.1 | 4.6 |
| Motala12 | (Lazaridis *et al.*, 2014) | GRCh38.p12 | 5500.0 | 7603.6 | 803.7 | 9.5 |

The runtime is given in seconds and the average runtime of ten independent runs. The runtime was measured for four bam files and includes reading the bam file, reconstruction of the mapped reads, and calculating and plotting the damage patterns. The factor is calculated by dividing the runtime of DamageProfiler by the runtime of mapDamage2.

### References


Benson,D.A. *et al.* (2013) GenBank. *Nucleic Acids Res.*, **41**, D36–42.

Gretzinger,J. *et al.* (2019) Large-scale mitogenomic analysis of the phylogeography of the Late Pleistocene cave bear. *Sci. Rep.*, **9**, 10700.

Herbig,A. *et al.* (2016) MALT: Fast alignment and analysis of metagenomic DNA sequence data applied to the Tyrolean Iceman. *bioRxiv*, 050559.

Jónsson,H. *et al.* (2013) mapDamage2.0: fast approximate Bayesian estimates of ancient DNA damage parameters. *Bioinformatics*, **29**, 1682–1684.

Kircher,M. (2012) Analysis of high-throughput ancient DNA sequencing data. *Methods Mol. Biol.*, **840**, 197–228.

Lazaridis,I. *et al.* (2014) Ancient human genomes suggest three ancestral populations for present-day Europeans. *Nature*, **513**, 409–413.

Li,H. (2013) Aligning sequence reads, clone sequences and assembly contigs with BWA-MEM. *arXiv [q-bio.GN]*.

Li,H. (2018) Minimap2: pairwise alignment for nucleotide sequences. *Bioinformatics*, **34**, 3094–3100.

Li,H. and Durbin,R. (2009) Fast and accurate short read alignment with Burrows-Wheeler transform. *Bioinformatics*, **25**, 1754–1760.

Peltzer,A. *et al.* (2016) EAGER: efficient ancient genome reconstruction. *Genome Biol.*, **17**, 60.

Quadrato,G. *et al.* (2017) Fast gapped-read alignment with Bowtie 2. *Nat. Methods*, **545**, 357–359.

Renaud,G. *et al.* (2017) gargammel: a sequence simulator for ancient DNA. *Bioinformatics*, **33**, 577–579.

Schubert,M. *et al.* (2016) AdapterRemoval v2: rapid adapter trimming, identification, and read merging. *BMC Res. Notes*, **9**, 88.

Schubert,M. *et al.* (2012) Improving ancient DNA read mapping against modern reference genomes. *BMC Genomics*, **13**, 178.

Schuenemann,V.J. *et al.* (2017) Ancient Egyptian mummy genomes suggest an increase of Sub-Saharan African ancestry in post-Roman periods. *Nat. Commun.*, **8**, 15694.

Schuenemann,V.J. *et al.* (2018) Ancient genomes reveal a high diversity of Mycobacterium leprae in medieval Europe. *PLoS Pathog.*, **14**, e1006997.

Vågene,Å.J. *et al.* (2018) Salmonella enterica genomes from victims of a major sixteenth-century epidemic in Mexico. *Nat Ecol Evol*, **2**, 520–528.

Warinner,C. *et al.* (2014) Pathogens and host immunity in the ancient human oral cavity. *Nat. Genet.*, **46**, 336–344.
